# Characterization of a COQ8A-ataxia mouse model with E548K single-site mutation: distinct and comparable findings relative to a loss-of-function mutation

**DOI:** 10.1101/2025.04.23.650169

**Authors:** Débora Farina Gonçalves, Natacha Broyer, Maïté Carret Pierrat, Emmanuelle Sarzi Campillo, Laurence Reutenauer, Hélène Puccio

## Abstract

COQ8A-ataxia, also known as autosomal recessive cerebellar ataxia type 2 (ARCA2), is a rare mitochondrial disorder caused by biallelic mutations in *COQ8A*, a gene encoding for a mitochondrial protein critical for coenzyme Q (CoQ) biosynthesis. Although there is no clear genotype-phenotype correlation in patients, loss-of function variants generally produce a cerebellar-restricted phenotype, while missense mutations are more frequently associated with multisystemic symptoms. The COQ8A^E551K^ variant has been reported at the homozygous state in individuals with early-onset disease and widespread systemic involvement. This study aimed to characterize the new *Coq8a*^*E548K*^ knocking mouse model, equivalent to the human E551K variant, and compare its phenotype to the complete *Coq8a*^*-/-*^ knockout mouse. Based on human data and preliminary data in the zebrafish, we hypothesized that the *Coq8a*^*E548K*^ knocking would present a more pronounced phenotype than the constitutive knockout. Contrary to our initial hypothesis, mice homozygous for the *Coq8a*^*E548K*^ allele exhibited no significant motor or cognitive impairments, nor muscle phenotype. Biochemically, the knocking *Coq8a*^*E548K*^ mutation led to variable instability of the COQ8A^E548K^ protein and reduced expression of proteins in the CoQ biosynthesis pathway, such as COQ5 and COQ7 in both cerebellum and muscle, similarly to the constitutive knockout. Despite this, mitochondrial function and tissue architecture remained intact, suggesting preserved cellular resilience. These findings highlight the complexity of genotype-phenotype correlations in COQ8A-related ataxia and provides a tool for investigating sub-threshold mitochondrial dysfunction.

## 1. Introduction

COQ8A-ataxia (former name : ARCA2, Autosomal recessive cerebellar ataxia type 2), is a rare recessive form of multisystemic ataxia. It is caused by biallelic mutations in *COQ8A* (former name: *ADCK3*), a gene encoding a mitochondrial protein involved in CoQ biosynthesis (1). CoQ is a redox-active lipid present in all membranes that is synthesized by a multiprotein complex, referred to as the CoQ synthome or complex Q, embedded in (2). Complex Q facilitates the enzymatic reactions necessary for synthesizing CoQ from its precursors (3). This involves serial reactions starting with addition of isoprenoid units and subsequent functional modifications in the benzoquinone ring to form the functional CoQ (4). COQ8A belongs to the UbiB family of atypical kinases (2). While its exact biochemical function remains undefined, COQ8A has been implicated in maintaining complex Q integrity and regulating CoQ biosynthesis (5). The current model is that COQ8A leverages its ATPase activity to support the formation of complex Q, potentially coupled with the extraction of CoQ lipids intermediates from the IMM to allow chemical modification of the benzoquinone head by the catalytic COQ proteins (6). Although variable, low CoQ levels have been identified in skeletal muscle and fibroblasts in some patients, displaying defects in mitochondrial homeostasis and signs of oxidative stress. While the pathophysiological mechanisms of COQ8A-ataxia are still being elucidated, emerging research, in particular from the constitutive mouse knockout, suggests mitochondrial dysfunction with impaired bioenergetics and mitochondrial oxidative stress, and Ca^+2^ imbalance as central features responsible for the specific neuronal vulnerability of the cerebellar Purkinje neurons (7).

Clinically, COQ8A-ataxia is characterized by slowly progressive early-onset cerebellar ataxia, combined with variable features including seizures or stroke-like episodes, cognitive decline, myopathy or exercise intolerance, and dystonia (8). The age of onset, clinical progression, and severity can vary considerably, often making diagnosis challenging (9). While no clear genotype-phenotype correlations have been established for COQ8A-ataxia, a recent study suggests that missense variants may be associated with more prevalent multisystemic involvement than biallelic loss-of-function variants (9,10). In contrast, patients harboring biallelic loss-of-function mutations often display phenotypes limited to cerebellar dysfunction. This distinction suggests that missense variants may exert dominant-negative or gain-of-function effects that contribute to broader clinical presentations. One such variant, *COQ8A*^*E551K*^, has been identified in homozygosity in a patient with early-onset ataxia (<2 years) and multisystemic symptoms including generalized tonic seizures and elevated lactate levels in cerebrospinal fluid at 2.5 years of age, progressive neurological deterioration, leading to inability to walk or speak by age 13 and the development of mild intellectual disability and epilepsy (11). Histological and biochemical studies of his skeletal muscle at 6 years demonstrated mitochondrial accumulation and accumulation of lipid droplets, as well as strong decrease in CoQ levels and loss of CII+CIII activity (11).

To investigate whether this mutation alone is sufficient to drive multisystemic pathology, we generated a knock-in model carrying the orthologous E548K mutation. We have previously generated a *Coq8a*^*-/-*^ constitutive knockout (KO) mouse model that recapitulates most hallmarks of COQ8A-ataxia, including a mild progressive cerebellar ataxia, exercise intolerance, increased epileptic susceptibility and mild memory impairment (5). In addition, *Coq8a*^*-/-*^ mice presented specific degeneration of Purkinje neurons (PNs), with significant gaps in the cerebellar PN layer and mitochondrial dysfunction, as evidenced by a marked reduction in the activities of Complex II and Complex IV (12). We aimed to determine whether E548K mutation results in a phenotype distinct or more severe than a complete COQ8A deficiency by comparing the *Coq8a* ^*E548K/E548K*^ mice with the established constitutive *Coq8a*^*-/-*^ knockout model (5). Our comprehensive analysis included behavioral, molecular, histological, and ultrastructural assessments. Contrary to our initial hypothesis, the *Coq8a* ^*E548K/E548K*^ mice did not present with any neurological phenotype, despite destabilization of complex Q.

## 2. Materials and Methods

### Mice generation and breeding

The E548K point mutation in the *Coq8a* gene was introduced into the mouse genome using CRISPR/Cas9-mediated gene editing in conjunction with homology-directed repair (HDR). To facilitate genotyping, a diagnostic RsaI restriction site was engineered into the repair template alongside the mutation. DNA from tail biopsies was amplified by PCR using the following primers:

- Forward: 5′-CCATCTGTAGGTGGCTCTCCTGGAC-3′
- Reverse: 5′-CACACCGGACCCCCAGGAGTAGCTG-3′

The resulting PCR products were digested with RsaI to distinguish between wild-type, heterozygous, and homozygous alleles based on the presence or absence of the restriction site. Digestion products were resolved by agarose gel electrophoresis to determine genotype.

The *Coq8a*^*E548K*^ heterozygous mice were bred to generate homozygous mutants, and the genotypes of offspring were determined. Mice were maintained in a pathogen-free animal facility with a 12-hour light/dark cycle and provided with standard rodent chow (D04, SAFE, Villemoisson-sur-Orge, France) and water ad libitum. All animal procedures were performed in accordance with institutional animal care guidelines and approved by the local ethical committee (APAFIS# 36059-2022021711458000).

### Behavioral phenotyping

Behavioral phenotyping was performed to evaluate motor coordination, balance, motor skills, neuromuscular function, gait, and short-term spatial memory in mice. To assess motor coordination and balance, the accelerating rotarod test (Panlab, Barcelona, Spain) was used as previously described (5). Mice were first habituated to the apparatus by undergoing three daily trials on the rod rotating at a constant speed of 4 rpm. Upon successful habituation, the testing phase began with the rod accelerating from 4 rpm to 40 rpm over a period of 5 minutes, over three consecutive days. Neuromuscular function and peripheral coordination were further assessed using the hanging wire test, as previously described (5). Mice were placed on the wire with their front limbs, and the time until they successfully placed their hind limbs onto the wire was recorded using a stopwatch. Mice were required to hold the wire with their hind limbs for at least 2 seconds to be considered a successful trial. The linear movement was evaluated by footprint analysis as previously described (13). We assessed linearity, defined as the change in angle between consecutive right-right steps, calculated by drawing a line perpendicular to the direction of travel. A high linearity score indicates more nonlinear movement. The work spatial memory was assessed by the Y-maze test, as reported before (14). Data were collected by tracking the time spent in each arm, with results reflecting short-term spatial memory based on the differential exploration of arms. The beam walking task was performed as previously described (8). Three trials were performed per mouse, recording the time taken to cross and the number of foot slips (12). Data were analyzed to assess motor coordination and balance, with statistical comparisons between groups.

### Antibodies and Western Blots

Western blots were performed according to standard protocol (12), including separation of proteins by SDS Tris-Glycine PAGE. Antibodies were diluted as follows: anti-COQ8A (1/1000 generated in-house against the peptide KQMTKTLNSDLGPHWRDKC), anti-COQ8B (1/1000 generated in-house against the peptide PGGSLQHEGVSGLGC), anti-COQ7 (1/1000, Santa cruz), anti-COQ5 (Proteintech 17453-1-AP, 1/1000).

### Hematoxylin Phloxine coloration

Ctrl and *Coq8a* ^*E548K/E548K*^ (30 weeks) were euthanized by cervical dislocation. The cerebellum and quadriceps muscles tissues were embedded in OCT and sectioned at 10 µm thickness. Sagittal sections were stained using a Hematoxylin-Phloxine protocol. Sections were hydrated, stained in Gill’s hematoxylin for 10 minutes, differentiated in acid alcohol, and counterstained with 0.1% Phloxine B for 5 minutes. Sections were dehydrated through ethanol, cleared in xylene, and mounted with a cover slip. Imaging was performed using a slide sanner Zeiss axio Z1 microscope,identifying Purkinje neurons and quadriceps muscle fibers. Quantification of PN number was conducted using ZEISS ZEN lite® software and statistical analysis was performed with GraphPad Prism.

### Immunofluorescence

Whole brains from mice were harvested and fixed in 4% paraformaldehyde (PFA) overnight (ON). The brains were then cryoprotected in 30% sucrose in PBS for 48 hours, snap-frozen in isopentane chilled on dry ice, and embedded in OCT. Cerebellum saggital sections of 30 µm were cut using a cryostat. A floating immunohistochemistry (IHC) protocol was followed, as previously described (15). Primary antibodies against anti-COQ8A (generated in-house against the peptide KQMTKTLNSDLGPHWRDKC), anti-COQ7 and anti-COQ5 (Proteintech 17453-1-AP, 1/1000) were used at a concentration of 1:250.

### Histoenzymatic staining of mitochondria complexes II and IV

Cerebella from 30-week-old mice were cryoprotected, embedded in OCT, and sectioned at 14 µm using a cryostat. The sections were incubated at 37°C for 40 minutes in freshly prepared histo-enzymatic media for each complex. For Complex II, the solution contained 1.23 mg/ml NBT, 35.12 mg/ml sodium succinate (Sigma, S2378), 0.0613 mg/ml Phenazine methosulfate (PMS; Sigma, P9625), and 0.065 mg/ml sodium azide (Sigma, S8032) in PBS. For Complex IV, the solution consisted of 0.5 mg/ml 3,3′-diaminobenzidine Tetrahydrochloride (DAB; Sigma, D7304), 1 mg/ml cytochrome c (Sigma, C2506), and 2 μg/ml bovine catalase (Sigma, C9322) in PBS (pH 7.4) (16). Imaging was performed using a slide sanner Zeiss axio Z1 microscope, identifying Purkinje neurons and muscle structures. Quantification was conducted using ZEISS ZEN lite® software and statistical analysis was performed with GraphPad Prism.

### Electron Microscopy

Animals were intracardiac perfused with PBS solution and following the fixation started with a solution of 2.5% glutaraldehyde and 2.5% PFA diluted in PBS. Tissues were kept in 2.5% glutaraldehyde + 2.5% PFA solution. For the process of post fixation tissues were rinsed in 1% osmium tetroxide-PBS (2 hr, 4 °C), dehydrated, and embedded in Epon. Regions of interest were localized on 2 μm and 5um sections and stained with toluidine blue. Ultrathin sections from selected areas were stained with uranyl acetate and lead citrate and examined with a Philips 208 electron microscope, operating at 80 kV (5).

### Statistical analysis

Graphs and analysis were made using Prisma 9.0. Statistical analysis was performed by a two-way ANOVA for behavioral tests, as the animals were evaluated at multiple time points throughout their life. For all other comparisons, a Student’s t-test was employed to assess differences between the control and Coq8a^E548K/E548K^ groups.

## 3. Results

### COQ8A^E548K^ mutation at homozygosity in mice does not induce motor deficit

We generated a transgenic mouse line carrying a missense mutation that substitutes glutamate for lysine at position 548 (E548K), corresponding to the human COQ8A^E551K^ variant using CRISPR-Cas9 strategy (see methods). To evaluate the impact of the mutation, we conducted a longitudinal behavioral analysis on *Coq8a*^*E548K/E548K*^ mice from 5 and 30 weeks of age. Unlike *Coq8a*^*-/-*^ mice which exibit progressive motor deficit starting at 10 weeks of age (5), *Coq8a*^*E548K/E548K*^ mice displayed largely normal motor coordination. No significant differences were observed in rotarod performance (Fig. 1A and 1D), footprint linearity (Fig. 1C and 1F), or missteps on the bar test (Fig. 1A supplementary data) compared to control. While phenotype variability was noted across time points and individuals, the homozygous E548K mutation did not lead to over behavioral impairments. Y-maze testing also showed no significant differences in exploratory behavior, suggesting preserved short-term spatial memory (Fig. 1B supplementary data). These findings support that the E548K variant does not results in substantial motor or cognitive decline under standard housing conditions.

**Figure 1:**
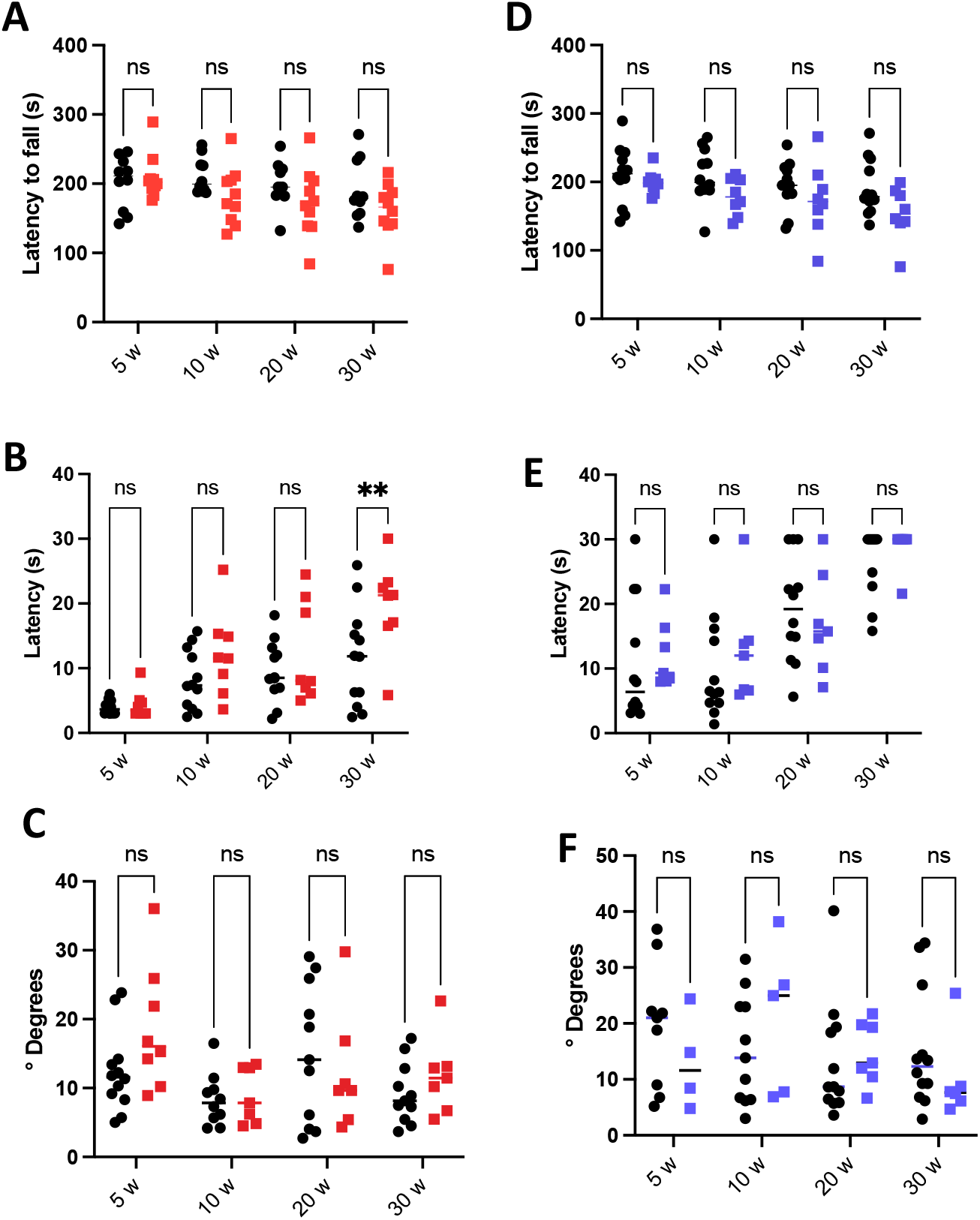
Coq8a^E548K/E548K^ mice do not exhibit alterations in motor coordination parameters throughout their lifespan. Behavioral performance of female (red squares) and male (blue squares) mice was compared with respective control groups (black dots). The tasks were performed at the following time points: 5 weeks, 10 weeks, 20 weeks, and 30 weeks. **(A, D)** Latency to fall in the reversed rotarod test. **(B, E)** Latency to regain balance in the hanging wire test. **(C, F)** Degree of linearity in the footprint test. Results are shown as individual values. Statistical analysis was performed using two-way ANOVA (**p < 0.01);

### COQ8A^E548K^ mutation disturbs complex Q in the cerebellum without affecting mitochondrial function of Purkinje neurons

We next sought to evaluate the impact of the E548K missense mutation at the molecular level both on COQ8A and Complex Q stability. Western blot analysis revealed a reduction of COQ8A protein levels in cerebellum in *Coq8a*^*E548K/E548K*^ mice, with variability observed across individual animals (Fig. 2A), although mRNA expression remained stable (Fig. 2B).. These findings suggest that the E548K variant may destabilize COQ8A, leading to decreased protein levels. Interestingly, despite the disruption of COQ8A in the cerebellum, COQ8B, the very close paralog of COQ8A, did not show any significant changes in either gene expression (Supp Fig. 2A) or protein levels in the same tissue (Fig. 2C), suggesting the absence of compensation similarly to Coq8a^-/-^ mice. The COQ8A^E548K^ variant also led to a significant decrease in the levels of COQ5 and COQ7 proteins in the cerebellum (Fig. 2D and 2E). These findings suggest that the low levels of COQ8A^E548K^ impairs the steady state stability of CoQ complex proteins, as was previously observed in the Coq8a^-/-^ mice. In contrast to the effects observed following complete COQ8A loss, no the COQ8A^E548K^ variant did not affect PN morphology and survival (Fig. 2F), mitochondrial activity, with normal activities of Complex II (Fig. 2G) or Complex IV (Fig. 2H). Additionally, while COQ8A loss led to ultrastructural damage in Purkinje neurons, including dilated and fragmented Golgi apparatus and expanded cisternae of ribosome-rich endoplasmic reticulum in Coq8a^-/-^ mice (5), the COQ8A^E548K^ variant did not induce any ultrastructural alterations (Fig. 2I). Surprisingly, although we observed by western blot a decrease in COQ8A^E548K^ protein, immunofluorescence on PN did not show any decrease in COQ8A^E548K^ protein (Fig. 2J).

**Figure 2:**
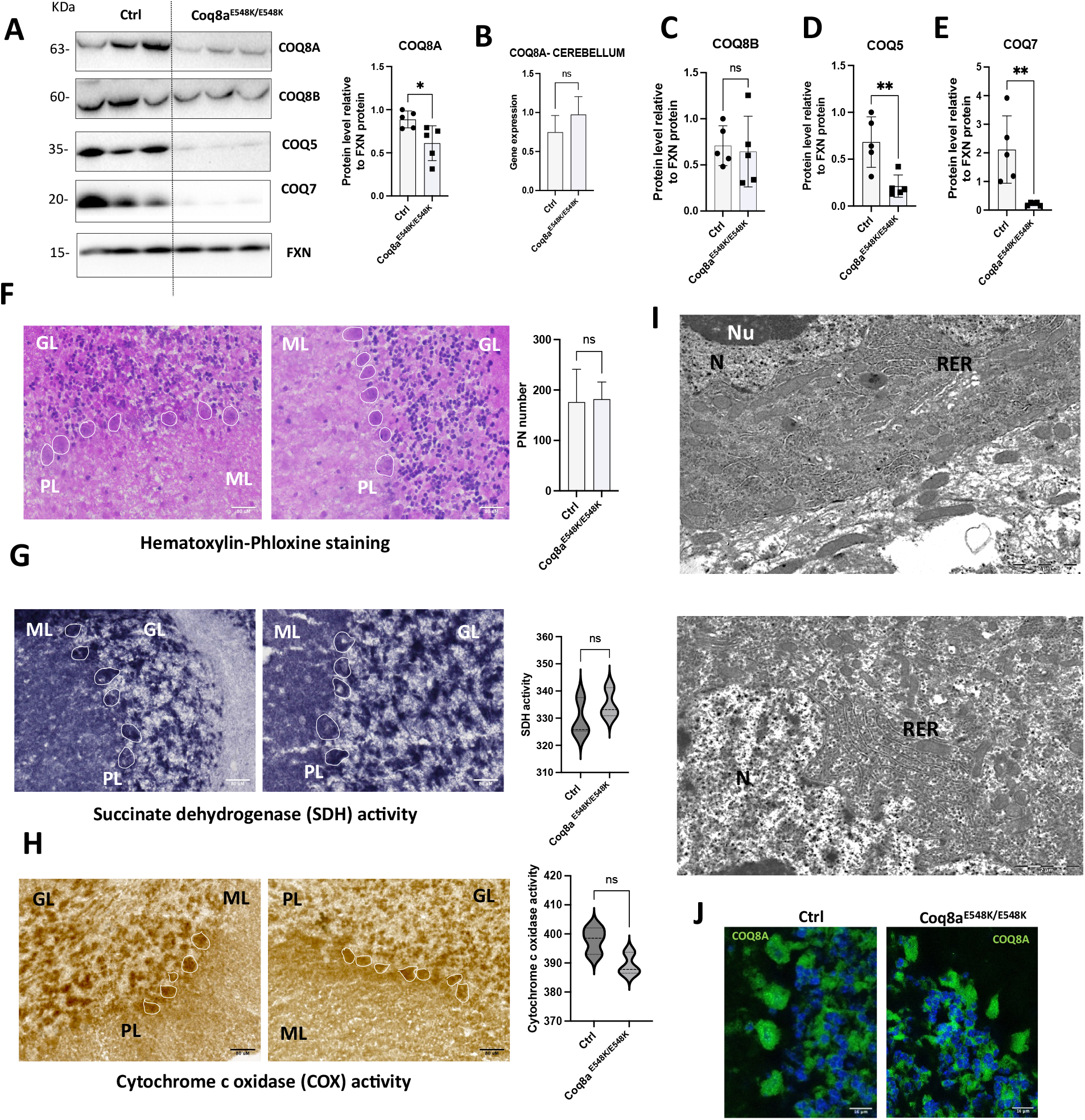
Coq8a^E548K/E548K^ mice exhibited dysregulation of the COQ synthome in the cerebellum, although mitochondrial and membrane structures of PN remained unaltered. **(A)** Representative western blots showing COQ protein levels (COQ8A, COQ8B, COQ5, COQ7) **(B)** COQ8A gene expression in the cerebellum. **(C-E)** Quantification of COQ synthome proteins in the cerebellum by western blot analysis. **(F**) Hematoxylin and phloxine staining in sagittal cerebellar sections, with Purkinje neurons **(PN)** highlighted in white. Scale bar = 80 µm. **(G, H)** SDH and COX histoenzymatic staining in sagittal cerebellar sections, with PN highlighted in white (n > 50 PN, at least 3 biological replicates). Scale bar = 80 µm. **(I)** Electron microscopy analysis of cerebellum showing rough endoplasmic reticulum (RER), nucleus (N), and nucleolus (Nu). Scale bar = 2 µm. **(J)** COQ8A immunostaining in Purkinje neurons. Scale bar = 16 µm. Data were obtained from *Coq8a*^E548K/E548K^ mice at 30 weeks of age. Quantification includes both male and female mice in each group (n = 3 to 5 animals per group). Results are presented as mean ± SD. Statistical significance was assessed using Student’s t-test (*p < 0.05; **p < 0.01); ns, not significant. 13

Additionally, other components of the COQ complex, such as COQ5 and COQ7, remain expressed in *Coq8a*^*E548K/E548K*^ PN cells (Fig. 3A supplementary data). The continued expression of COQ8A, along with other complex components, may be sufficient to sustain mitochondrial function specifically in those neurons, even in the presence of the mutation.

### COQ8A E548K Mutation Disturbs CoQ-Synthome in Muscle without Affecting Mitochondrial Function

In the quadriceps, COQ8A^E548K^ levels were also reduced, with expression levels varying across individual animals, while gene expression remained unaltered (Fig. 3A-C). Similar to the cerebellum, there was no compensatory increase in the expression of the paralog COQ8B (Fig. 3D). COQ5 and COQ7 proteins were significantly diminished (Fig. 3E-F), suggesting disruption of Complex Q. However, histological analysis showed normal muscle fiber morphology, and no increase in central nuclei was detected (Fig 3G), a hallmark of muscular degeneration seen in *Coq8a*^*-/-*^ mice. Furthermore, mitochondrial function, assessed via complex IV activity in both soleus and gastrocnemius muscles, remained unaffected (Fig. 3H). TEM analysis corroborated these findings, revealing intact mitochondrial cristae and overall ultrastructural preservation of muscle tissue (Fig. 3I). These results further support the notion that the E548K mutation does not lead to significant physiological dysfunction in muscle despite molecular dysregulation.

**Figure 3:**
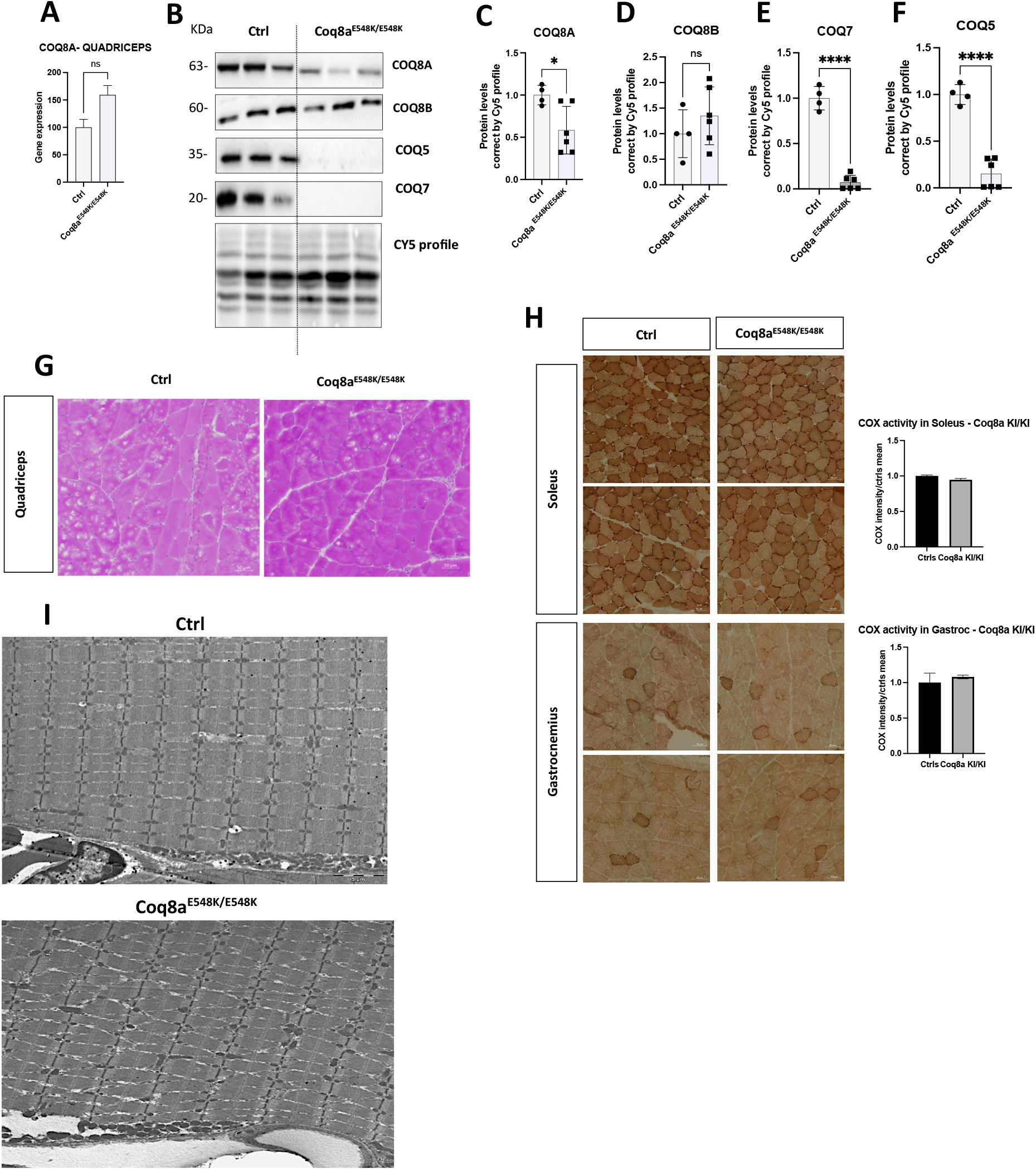
Coq8a^E548K/E548K^ mice presented derregulation of COQ synthome in quadriceps, but membrane structures and mitochonrial function are unaltered in muscle. **(A)** COQ8A gene expression in the quadriceps. **(B)** Representative western blots showing COQ protein levels (COQ8A, COQ8B, COQ5, COQ7). **(C-F)** Quantification of COQ synthome proteins in the quadriceps by western blot analysis. **(G**) Hematoxylin and phloxine staining in quadriceps sections. Scale bar = 50 µm. **(H)** COX histoenzymatic staining in soleus and gastrocnemius muscle sections. Scale bar = 50 µm. **(I)** Representative TEM images of quadriceps muscle tissue from control and Coq8a^E548K/E548K^ mice, showing no significant ultrastructural alterations between groups. Scale bar = 5 µm. Data were obtained from *Coq8a*^E548K/E548K^ mice at 30 weeks of age. Quantification includes both male and female mice in each group (n = 3 to 5 animals per group). Results are presented as mean ± SD. Statistical significance was assessed using Student’s t-test (*p < 0.05; ****p < 0.0001); ns, not significant.

## 4. Discussion

This study aimed to explore whether the COQ8A^E548K^ missense mutation, equivalent to the pathogenic human E551K variant, produced a more severe phenotype than COQ8A loss-of-function. *Coq8a*^*E548K/E548K*^ mice failed to recapitulate a multisystemic or worsened phenotype compared to *Coq8a*^*-/-*^ mice. Contrary to our initial hypothesis, the COQ8A^E548K^ variant did not lead to a multisystemic phenotype in mice. Our behavioral analyses revealed no significant deficits in coordination across several assays, in contrast to the progressive motor impairment seen in Coq8a-null mice. The retention of COQ8A expression in E548K mice, albeit at reduced levels, likely explains the milder phenotype. This residual protein presence appears sufficient to maintain CoQ biosynthetic function in high-demand tissues such as cerebellum and muscle. The variability in behavioral phenotypes suggests subtle modulation of neuronal function that does not reach a pathological threshold under standard conditions.

Biochemically, the E548K mutation induces dysregulation of the CoQ biosynthetic complex, with reduced expression of COQ5 and COQ7 in both cerebellum and muscle, resembling the molecular signature of complete COQ8A loss. Yet, unlike the knockout model, this dysregulation does not result in downstream mitochondrial dysfunction or overt structural degeneration. In Purkinje neurons, COQ8A, COQ5, and COQ7 remain detectable, and mitochondrial activity is preserved, highlighting the resilience of these cells in the context of partial CoQ biosynthesis disruption. It has been demonstrated that the COQ8A protein is essential for CoQ biosynthesis, and that various mutations can affect its activity. For example, the E551K mutation alters COQ8A’s structure, leading to thermal instability and electrostatic clashes (2). While these changes likely disrupt proper protein folding, they do not appear to completely abolish its function in *Coq8a*^*E548K/E548K*^ mice.

Our findings underscore the concept of a phenotypic threshold in mitochondrial diseases. As proposed in other mitochondrial disorders, cellular dysfunction becomes clinically relevant only when biochemical impairment surpasses a certain threshold (17). In this case, the residual COQ8A activity may maintain bioenergetic homeostasis below that threshold, preventing the manifestation of severe symptoms.

In conclusion, the COQ8A^E548K^ missense mutation results in partial disruption of the CoQ biosynthetic pathway. Although it alters the molecular landscape of CoQ metabolism, it does not significantly impair mitochondrial function or tissue integrity within the timeframe of this study. These findings suggest that E548K acts as a hypomorphic allele, enabling survival and relatively normal function despite detectable biochemical perturbations. Additionally, the fact that the homozygous E551K variant in humans leads to an early and multisystemic phenotype— characterized by seizures, generalized tonic-clonic episodes, and progressive neurological deterioration—highlights the importance of considering the complex interaction between genetic mutations, cellular processes, and environmental factors when developing therapeutic strategies.

## Supplementary DATA

**Figure 1:**
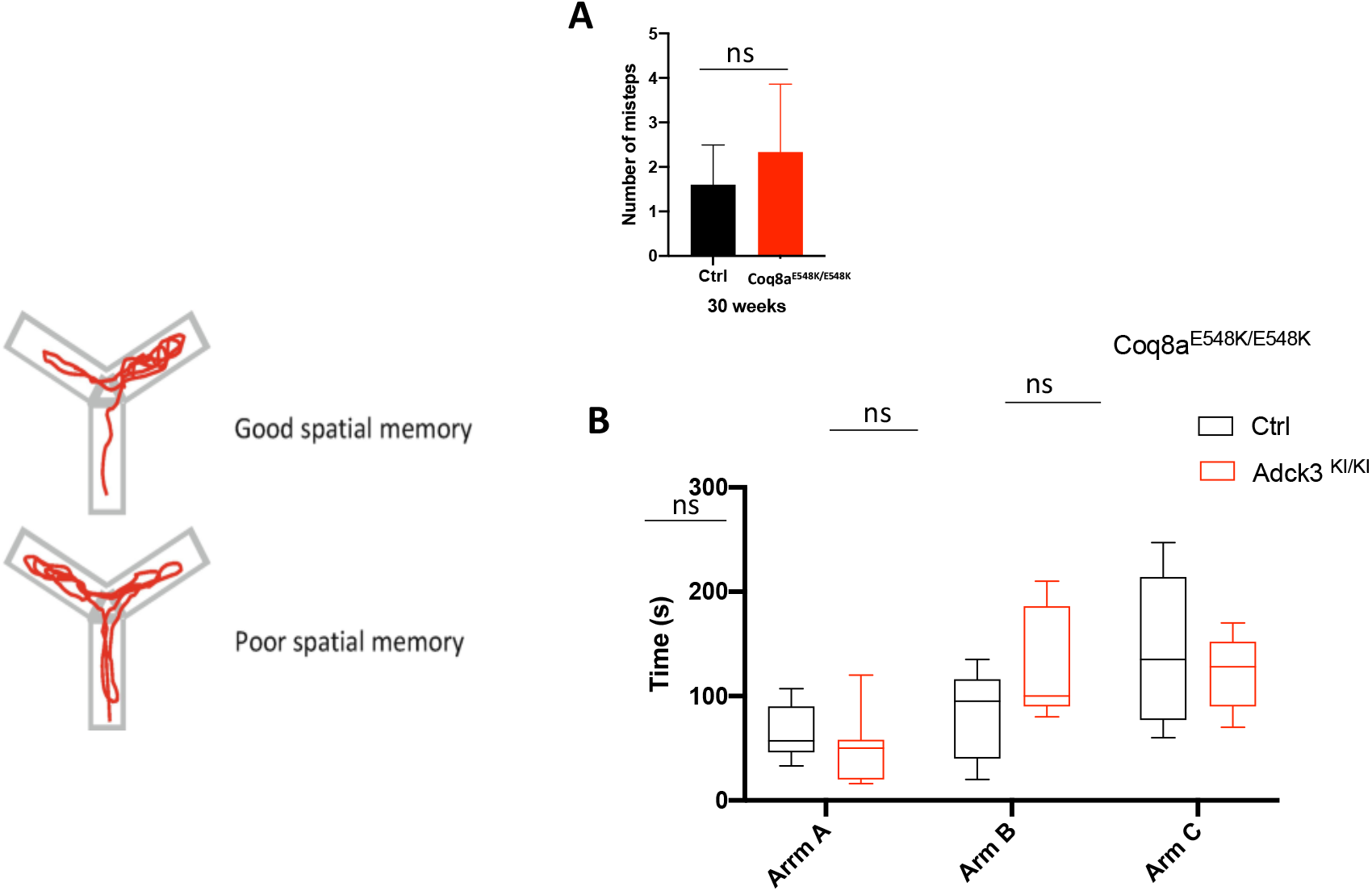
Coq8a ^E548K/E548K^ mice do not exhibit impairments in spatial memory or refined motor coordination. **(A)** Number of missteps in the bar test, evaluating motor coordination. Data were obtained from 30-week-old Coq8a ^E548K/E548K^ mice **(B)** Quantification of time spent exploring different arms in the Y-maze test, assessing spatial memory. (n = 9 to 11 animals per group). Results are presented as mean ± SD. Statistical significance was assessed using Student’s t-test comparing control and Coq8a ^E548K/E548K^ mice; ns, not significant.

**Figure 2:**
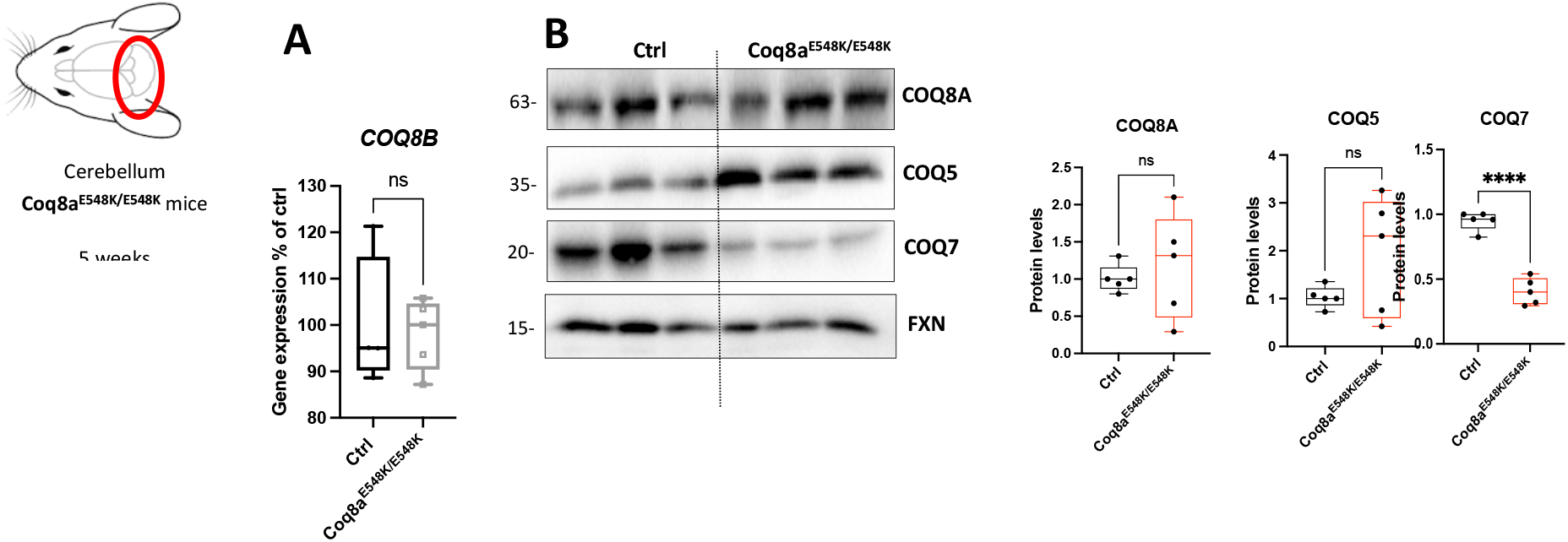
CoQ synthome in cerebellum of Coq8a^E548K/E548K^ mice at 5 weeks of age. (A) COQ8B gene expression Coq8a^E548K/E548K^ mice (B) Representative western blot analysis and quantification showing the expression of COQ8A, COQ5, and COQ7 proteins in the cerebellum of wild-type and Coq8a^E548K/E548K^ mice at 5 weeks. Results are expressed as mean ± SD. Statistical significance was determined using Student’s t-test (****p < 0.0001); ns, not significant.

**Figure 3:**
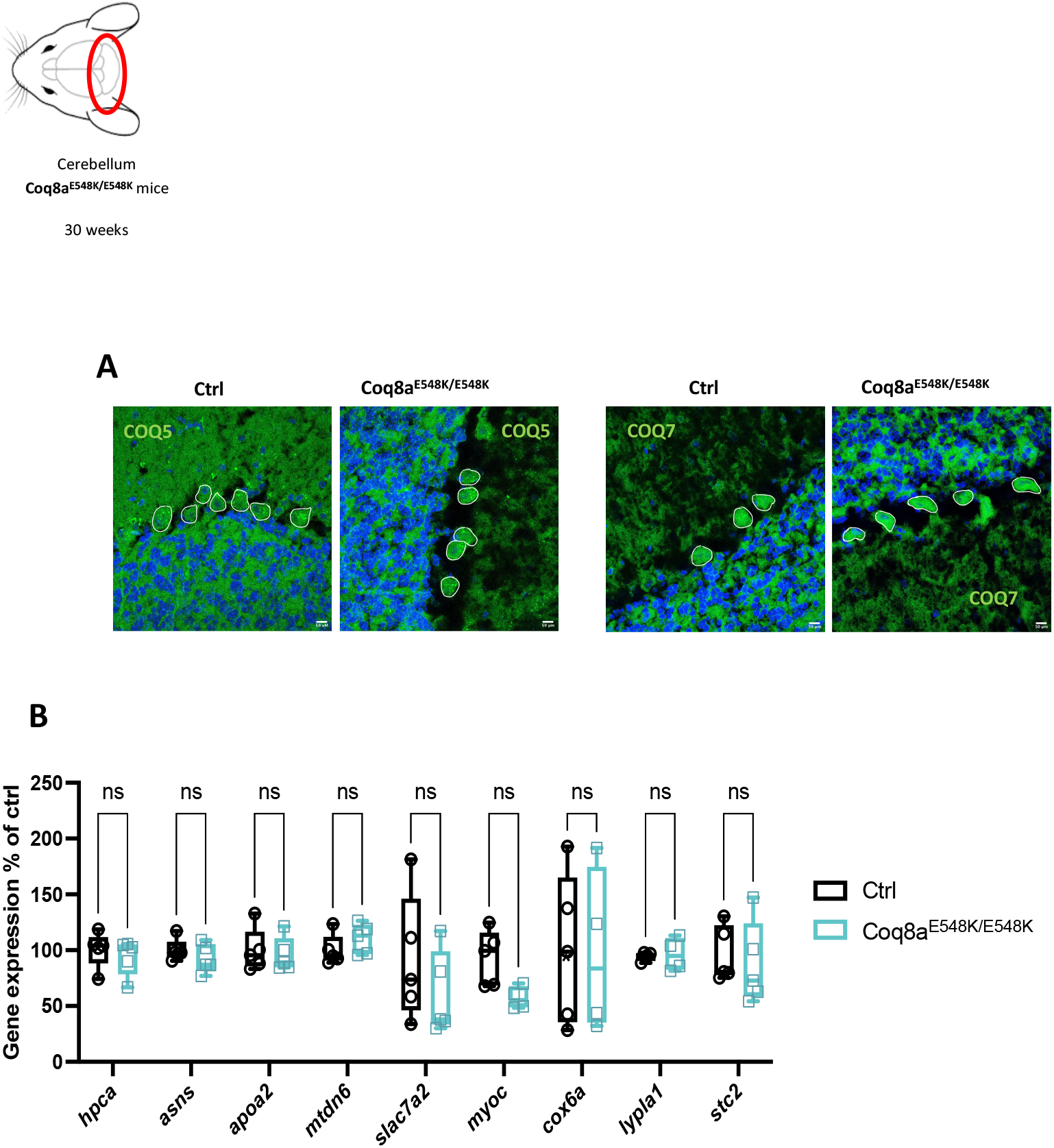
Expression of key genes in the cerebellum and COQ synthome in PN of Coq8a^E548K/E548K^ mice. **(A)** Immunofluorescence images showing the localization of COQ5 and COQ7 proteins in PN. **(B)** Gene expression analysis of selected genes in the cerebellum, identified based on their differential regulation in transcriptomic data from Coq8a^-/-^ mice. This analysis aimed to evaluate whether similar transcriptional changes would occur in Coq8a^E548K/E548K^ mice. Data were obtained from 30-week-old Coq8a^E548K/E548K^ mice. Results are presented as mean ± SD. Statistical significance was assessed using Student’s t-test; ns, not significant.

